# LimoRhyde: a flexible approach for differential analysis of rhythmic transcriptome data

**DOI:** 10.1101/283622

**Authors:** Jacob J. Hughey

**Author notes:** To whom all correspondence should be addressed.

## Abstract

As experiments to interrogate circadian rhythms increase in scale and complexity, methods to analyze the resulting data must keep pace. Although methods to detect rhythmicity in genome-scale data are well established, methods to detect changes in rhythmicity or in average expression between experimental conditions are often ad hoc. Here we present LimoRhyde (linear models for rhythmicity, design), a flexible approach for analyzing transcriptome data from circadian systems. Borrowing from cosinor regression, LimoRhyde decomposes circadian or zeitgeber time into multiple components, in order to fit a linear model to the expression of each gene. The linear model can accommodate any number of additional experimental variables, whether discrete or continuous, making it straightforward to detect differential rhythmicity and differential expression using state-of-the-art methods for analyzing microarray and RNA-seq data. In this approach, differential rhythmicity corresponds to a statistical interaction between an experimental variable and circadian time, whereas differential expression corresponds to the main effect of an experimental variable while accounting for circadian time. To demonstrate LimoRhyde’s versatility, we applied it to murine and human circadian transcriptome datasets acquired under various experimental designs. Our results show how LimoRhyde systematizes the analysis of such data, and suggest that LimoRhyde could become a valuable approach for assessing how circadian systems respond to genetic and environmental perturbations.

## Introduction

In diverse species from cyanobacteria to plants to mammals, circadian clocks drive rhythms in gene expression throughout the genome (Covington et al., 2008; Liu et al., 1995; Panda et al., 2002). Accordingly, transcriptome measurements have revealed circadian clocks’ influence on physiology, as well as potential applications for circadian medicine (Anafi et al., 2017; Hughey, 2017; Laing et al., 2017; Mure et al., 2018; Zhang et al., 2014). Transcriptome measurements are also beginning to reveal how circadian systems are affected by factors such as diet, infection, and cancer (Haspel et al., 2014; Masri et al., 2016; Tognini et al., 2017). The resulting datasets thus include samples not just from multiple time-points, but also from multiple conditions.

A common step in analyzing circadian or otherwise rhythmic transcriptome data is identifying which genes show evidence of rhythmic expression. This step can now be accomplished by various computational methods, including JTK_CYCLE and RAIN (Hughes et al., 2010; Hutchison et al., 2015; Thaben and Westermark, 2014). Importantly, though, these methods are only designed to detect rhythmic features (e.g., genes) based on samples from one condition. They are not designed to detect which features show a difference in rhythmicity between conditions (e.g., by comparing lists of rhythmic genes from each condition), and using them as such can lead to a high rate of false positives and false negatives (Thaben and Westermark, 2016). Indeed, the lack of a standard approach to analyze omics data from multiple conditions was highlighted in the recent guidelines for genome-scale analysis of biological rhythms (Hughes et al., 2017).

A classic approach for rhythm detection is cosinor regression (or harmonic regression), which is based on fitting a time series to the first harmonic of a Fourier series, i.e., sine and cosine curves of a set period (Cornelissen, 2014; Nelson et al., 1979). Because cosinor regression corresponds to a linear model, coefficients for even complex time series can be estimated efficiently using least squares. Inspired by cosinor regression, Thaben and Westermark recently made a significant advance in the statistically rigorous analysis of rhythmic transcriptome data from multiple conditions (Thaben and Westermark, 2016). Their method, called DODR, detects changes in rhythm amplitude, phase, and signal-to-noise ratio, which they call “differential rhythmicity.” DODR is relatively narrow in scope, though, as it is only designed to detect differential rhythmicity between two conditions. DODR cannot detect changes in average expression level between conditions, and cannot handle more complex experimental designs (e.g., with continuous variables such as age).

In addition to being a fundamental part of cosinor regression, linear models are one of two features shared by nearly all state-of-the-art methods for assessing differential expression in transcriptome data. The second is called empirical Bayes. While linear models provide the ability to handle complex experimental designs, empirical Bayes shares information across genes in order to make more stable estimates of gene-wise variance and thereby improve statistical power and accuracy (Smyth, 2004). These methods can also appropriately deal with read counts from RNA-seq (Soneson and Delorenzi, 2013). Despite these methods’ flexibility and widespread success, their application to circadian transcriptome data has been relatively limited (Hsu and Harmer, 2012; Montagner et al., 2016; Pembroke et al., 2015; Spörl et al., 2012). To our knowledge, there has been no unification of cosinor-based approaches with these state-of-the-art tools for differential expression.

We sought to develop a general approach to systematically analyze circadian transcriptome data from various experimental designs. Our approach, which we call LimoRhyde (linear models for rhythmicity, design), builds on cosinor regression to express complex circadian experiments in terms of a linear model, which makes circadian transcriptome data amenable to analysis by existing tools for differential expression. We validated our approach in the two-condition scenario by comparing it to DODR on six datasets from mice. To explore LimoRhyde’s flexibility, we then applied it to two datasets from humans. Our results suggest that LimoRhyde offers a valuable framework for assessing how rhythmic biological systems respond to genetic and environmental perturbations.

## Materials and Methods

All data and code to reproduce this study are available at https://figshare.com/s/31dcb1346ef7f4268aa6. The LimoRhyde R package is available at https://github.com/hugheylab/limorhyde.

### Processing the gene expression data

For the RNA-seq datasets (GSE73552 and E-MTAB-3428), we downloaded the raw reads, then quantified gene-level abundances (based on Ensembl Gene IDs) in units of transcripts per million (TPM) using salmon v0.8.2 and tximport v1.6.0 (Patro et al., 2017; Soneson et al., 2015). We kept for analysis only those genes having TPM ≥ 0.5 in at least half the samples. For all analyses and plots, we converted expression values to log_2_(TPM+1). For the microarray datasets, we downloaded the raw (Affymetrix) or processed (Agilent or Illumina) expression data from NCBI GEO, then used metapredict v0.0.0.9019 for mapping probes to Entrez Gene IDs, intra-study normalization, and log-transformation (Hughey and Butte, 2015). Details of all datasets are in Suppl. Table S1.

### Detecting rhythmic, differentially rhythmic, and differentially expressed genes using LimoRhyde and limma

To make circadian transcriptome data amenable to analysis using linear models, LimoRhyde follows the strategy of cosinor regression, decomposing zeitgeber or circadian time into a sine and cosine of period 24 h. Although this decomposition is the simplest, one could also decompose time based on multiple harmonics of the Fourier series or on periodic splines. Thus a single variable becomes at least two variables in the linear model. For data derived from several cycles in constant conditions, one could also include a linear time (e.g., time in free-run) to control for drift. Additional terms for condition, subject, or other covariates can be included as appropriate. In this approach, differential rhythmicity corresponds to a statistical interaction between the experimental factor of interest (e.g., genotype) and each term related to zeitgeber/circadian time. Differential expression, meanwhile, corresponds to the main effect of the experimental factor of interest.

After constructing the linear model, the transcriptome data can be analyzed using multiple existing methods based on linear models and empirical Bayes. In this paper, we used limma v3.34.9 (Ritchie et al., 2015; Smyth, 2004). For all datasets (microarray and RNA-seq), we used limma with largely the default settings, expect we allowed it to fit a mean-variance trend across genes (limma-trend) (Law et al., 2014). To control the false discovery rate at every step of the analysis, p-values were converted to q-values using the method of Benjamini and Hochberg (Benjamini and Hochberg, 1995).

In the datasets from mice, which have discrete time-points spaced throughout the circadian cycle, we detected genes with rhythmic expression using RAIN (see next section). In the dataset based on samples from human brain (GSE71620), one experimental factor (age) is continuous and the zeitgeber time-points are approximately randomly distributed. Therefore, to calculate a q-value of rhythmicity (accounting for age), we first used LimoRhyde to construct an additive model with terms for age, brain region, and zeitgeber time. The model does not include a term for donor, because although each donor has a corresponding sample from each of two brain regions, those two samples correspond to the same age and the same zeitgeber time, making it impossible to reliably account for inter-donor variation. We then used limma to perform a moderated F-test on the coefficients corresponding to the two terms for zeitgeber time.

### Detecting rhythmic genes using RAIN

For datasets with two conditions, our goal was to detect genes rhythmic in at least one condition. For datasets with discrete time-points (all mouse datasets), we followed a similar procedure as used previously (Thaben and Westermark, 2016). We first ran RAIN v1.12.0 (default settings and period 24 h) separately on the samples from each condition, which resulted in a p-value for each gene in each condition. We then used the minimum p-value for each gene to calculate q-values of rhythmicity (q_rhy_). Comparing acrophase across conditions only makes sense, if the gene is rhythmic in both conditions. We calculated q-values of being rhythmic in both conditions (q_rhy,max_) similarly, but using the maximum p-value instead of the minimum.

### Detecting differentially rhythmic genes using DODR

We used DODR v0.99.2 with the default settings (Thaben and Westermark, 2016), which performs both robustDODR and robustHarmScaleTest. We used only the former, which tests for a combination of amplitude and phase change and corresponds to the moderated F-test in limma. The robustHarmScaleTest, which tests for a difference in noise level of rhythmic expression, was less informative in our experience. If desired, one could test for differential variability of rhythmic expression in the LimoRhyde/limma framework using a method called DiffVar (Phipson and Oshlack, 2014).

### Comparing LimoRhyde and DODR for detecting differential rhythmicity

To evaluate the agreement between LimoRhyde (followed by limma) and DODR in calling genes differentially rhythmic, we calculated Cohen’s kappa at various q-value cutoffs using irr R package v0.84. To estimate each method’s tendency to call false positives in each dataset, we first identified genes rhythmic in at least one condition using RAIN on the true sample labels, then permuted the sample labels (wild-type or knockout) within samples from the same time-point. This strategy attempts to preserve rhythmic expression patterns, but remove differential rhythmicity. For each dataset, we then calculated the mean number of differentially rhythmic genes (across 50 permutations) at various q-value cutoffs. Because the order of magnitude of differentially rhythmic genes varies across datasets, we summarized the overall results using the geometric mean.

To ensure a sufficient number of rhythmic genes for comparison, we used a cutoff of q_rhy_ ≤ 0.1 for five of the six datasets. We used a cutoff of q_rhy_ ≤ 0.15 for E-MTAB-3428, which has only four time-points and eight samples per genotype.

### Calculating gene-wise rhythmic parameters using ZeitZeiger

We estimated rhythm amplitude and zeitgeber/circadian time of peak expression (acrophase) using ZeitZeiger v1.0.0.5 with default settings (Hughey et al., 2016; Hughey and Butte, 2016). For the mouse datasets, we ran ZeitZeiger separately on the samples from each condition. For the human brain dataset, to calculate each gene’s overall rhythmic properties, we used LimoRhyde and limma to adjust the expression values for age and brain region, then ran ZeitZeiger on the residual expression values from all samples. To estimate the change in rhythmic properties with age, we split donors into a younger 50% and older 50%, then calculated the rhythm amplitude and acrophase on the unadjusted expression values within each cohort. For all datasets, we calculated Δamplitude as the arithmetic difference in rhythm amplitude, and Δacrophase as the circular difference (constrained between −12 and +12 h).

### Performing gene set analysis using CAMERA

The CAMERA method (Wu and Smyth, 2012) is part of the limma R package. To identify gene sets enriched for differential expression, we used the “camera” function, which takes as input an expression matrix, a list of gene sets, a design matrix corresponding to a linear model, and a single contrast (e.g., genotype or age). As in the limma analysis, we allowed camera to fit a mean-variance trend. To identify gene sets enriched for differential rhythm amplitude, we used the “cameraPR” function (default settings), which takes as input a vector of gene-wise statistics (in our case, Δamplitude) and a list of gene sets. We used the mouse and human C5 GO (gene ontology) gene sets of MSigDB v5.2 (Liberzon et al., 2011), which are available at http://bioinf.wehi.edu.au/software/MSigDB/index.html. The gene sets are based on Entrez Gene IDs, so for the gene set analysis of GSE73552, we mapped Ensembl Gene IDs to Entrez Gene IDs using the org.Mm.eg.db R package, keeping only genes with a one-to-one mapping.

## Results

### Applying LimoRhyde to circadian transcriptome data from a two-condition design

To develop a workflow for using LimoRhyde, we first sought to analyze a circadian transcriptome dataset that is representative of a common experimental design, in which samples are acquired at discrete time-points throughout the circadian cycle in two conditions. We selected a dataset that included samples taken every 4 h from livers of wild-type and Arntl-/-mice under night-restricted feeding in LD 12:12, with gene expression measured by RNA-seq (Atger et al., 2015). Starting with the RNA-seq reads, we estimated gene-level abundances using salmon and tximport (Patro et al., 2017; Soneson et al., 2015). Using RAIN (Thaben and Westermark, 2014), we then identified 2,434 genes rhythmic in at least one genotype (q_rhy_ ≤ 0.01; Fig. 1A and Suppl. Fig. S1A).

**Figure 1.**
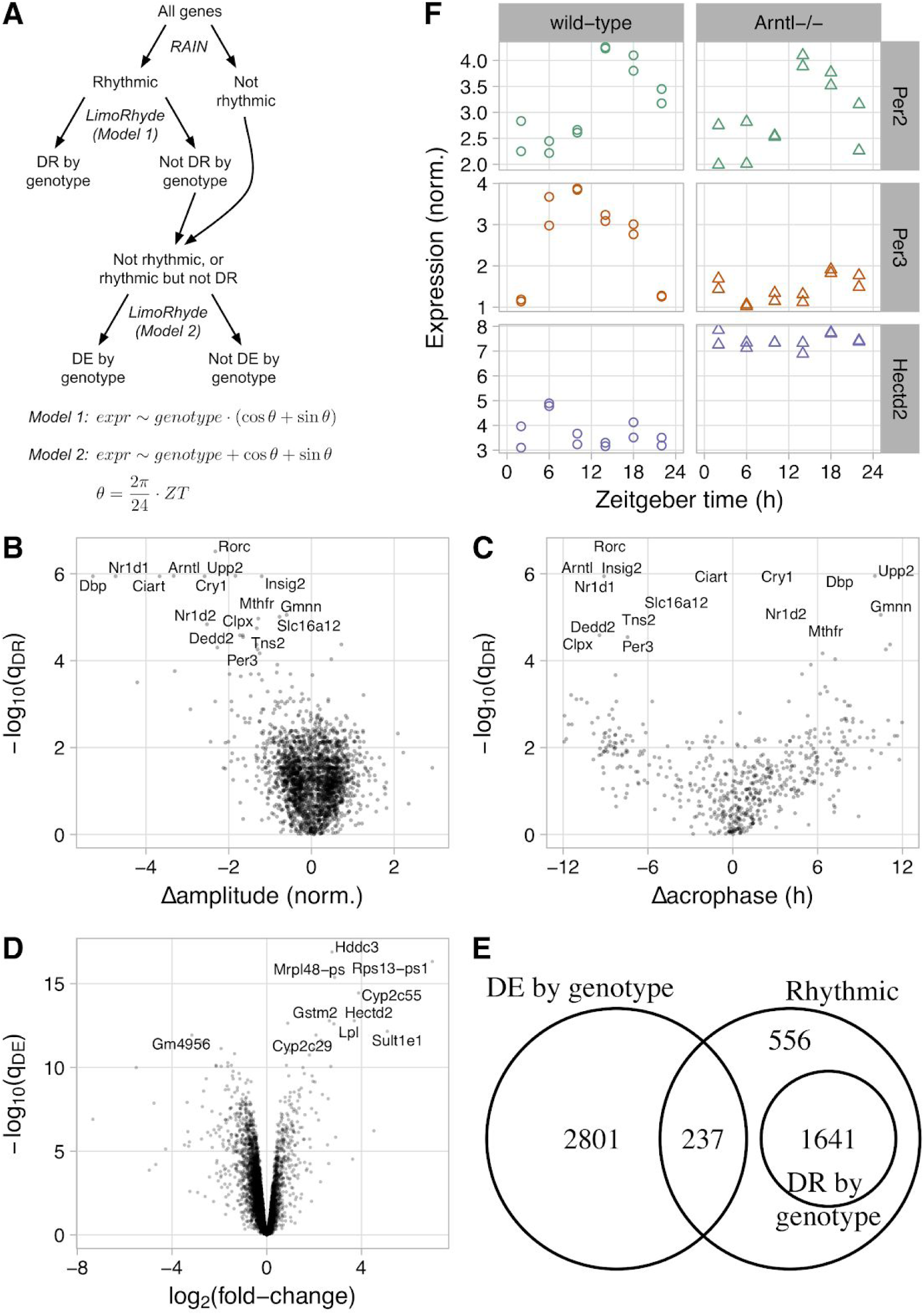
Using LimoRhyde to analyze circadian transcriptome data from livers of wild-type and Arntl-/-mice under night-restricted feeding (GSE73552). **(A)** Schematic of workflow and linear model formulae used to detect rhythmic, differentially rhythmic (DR), and differentially expressed (DE) genes. ZT corresponds to zeitgeber time. Each formula corresponds to a linear regression fit, in which expression of each gene (expr) is modeled as a function of the variables to the right of the the tilde. The formulae do not show intercepts or coefficients. Multiplication in a model formula indicates inclusion of the main effects and the interaction. **(B)** Scatterplot of −log_10_(q_DR_) vs. Δamplitude. q_DR_ corresponds to a rhythmic gene’s q-value of differential rhythmicity, calculated using LimoRhyde and limma (Model 1). Δamplitude corresponds to the change in rhythm amplitude between genotypes, where a negative value indicates lower amplitude in Arntl-/-. Only genes with a q_rhy_ ≤ 0.01, where q_rhy_ is the q-value of rhythmicity, were considered. In (B), (C), and (D), each point represents a gene. In (B) and (C), the 16 rhythmic genes with the highest −log_10_(q_DR_) are labeled. **(C)** Scatterplot of −log_10_(q_DR_) vs. Δacrophase. The latter corresponds to the change in zeitgeber time of peak expression, where a positive value indicates a phase advance in Arntl-/-. Only genes with q-value for rhythmicity in both genotypes ≤ 0.2 are shown. **(D)** Scatterplot of −log_10_(q_DE_) vs. log_2_ fold-change, both calculated using LimoRhyde and limma (Model 2). q_DE_ corresponds to the q-value of differential expression. A positive log_2_ fold-change indicates higher average expression in Arntl-/-. The 10 genes with the highest −log_10_(q_DE_) are labeled. **(E)** Venn diagram of genes meeting criteria for rhythmicity (q_rhy_ ≤ 0.01), differential rhythmicity (q_DR_ ≤ 0.1), and differential expression (q_DE_ ≤ 0.1). **(F)** Plots of three example genes. Each point represents a sample. Based on the criteria, Per2 is classified as rhythmic only, Per3 as differentially rhythmic, and Hectd2 as differentially expressed only.

We next used LimoRhyde to express the experimental design in terms of a linear model (Fig. 1A), in order to use a method called limma to determine which rhythmic genes showed evidence of differential rhythmicity between wild-type and Arntl-/-mice. Limma is a general method for analyzing microarray and RNA-seq data based on linear models and empirical Bayes (Ritchie et al., 2015; Smyth, 2004). We used limma to calculate a moderated F-statistic for each rhythmic gene, which tests the null hypothesis that both coefficients corresponding to the interaction between genotype and zeitgeber time are zero. This amounts to testing for a difference in a combination of rhythm amplitude and phase (Thaben and Westermark, 2016). Of 2,434 rhythmic genes, 1,641 genes were differentially rhythmic at a cutoff of q_DR_ ≤ 0.1 (Suppl. Fig. S1B). Of the 16 genes with the lowest q_DR_, 8 genes are part of or directly driven by the core circadian clock (Rorc, Arntl, Nr1d1, Dbp, Cry1, Ciart, Nr1d2, and Per3).

Although the moderated F-statistic can provide evidence of a change in rhythmicity, it does not indicate the nature of the change. Therefore, for each rhythmic gene, we used ZeitZeiger (Hughey et al., 2016) to quantify the rhythm amplitude and the zeitgeber time of peak expression (acrophase) in wild-type and Arntl-/-mice. We found that the genes with the strongest evidence for differential rhythmicity had strongly reduced rhythm amplitude in Arntl-/-mice (Fig. 1B). Among genes that exhibited at least moderate evidence of rhythmicity in each genotype (q_rhy,max_ ≤ 0.2; see Materials and Methods), changes in acrophase were widely distributed (Fig. 1C). As expected (Thaben and Westermark, 2016), genes with stronger evidence of differential rhythmicity tended to have larger absolute changes in acrophase.

We then used a simpler linear model, one lacking an interaction between genotype and zeitgeber time, to identify genes differentially expressed between wild-type and Arntl-/-mice. Here differential expression refers to a difference in average expression level between genotypes, accounting for possible rhythmicity. For this step, we considered only the 11,737 genes for which there was not strong evidence of differential rhythmicity (q_DR_ > 0.1) or for which differential rhythmicity was not examined (q_rhy_ > 0.01). Among these genes, 3,038 genes were differentially expressed (q_DE_ ≤ 0.01), of which 301 genes had an absolute log_2_ fold-change > 1 (Fig. 1D).

Finally, to complement the gene-wise analysis, we used a method called CAMERA to perform gene set analysis (Wu and Smyth, 2012). Because methods such as CAMERA are unable to work directly with the F-statistics of differential rhythmicity (which have only a positive sign), we instead plugged into CAMERA the differences in rhythm amplitude as quantified by ZeitZeiger. Consistent with the gene-wise analysis, four of the five top-ranked gene sets with altered rhythm amplitude were related to circadian rhythms (all with q ≤ 10^−8^ and reduced amplitude in Arntl-/-; Suppl. Table S2). Gene sets enriched for differential expression, meanwhile, tended to be related to the ribosome and various catabolic processes (with increased expression in Arntl-/-; Suppl. Table S3).

Given criteria for rhythmicity, differential rhythmicity, and differential expression, the assignment of genes to each group can be expressed as a Venn diagram (Fig. 1E). To illustrate the various expression patterns, we show three genes as examples (Fig. 1F). Taken together, these results suggest that LimoRhyde provides a cohesive framework for differential analysis of circadian transcriptome data.

### Comparing LimoRhyde and DODR in the assessment of differential rhythmicity

The recently developed method DODR is designed to detect differential rhythmicity between two conditions. As in LimoRhyde, differential rhythmicity in DODR is defined as a statistical interaction in a linear model based on cosinor regression. Although DODR does not use empirical Bayes to share information between genes, it does use rank-based statistics to achieve robustness to outlier samples.

To compare LimoRhyde (followed by limma) and DODR, we assembled six circadian transcriptome datasets (four microarray, two RNA-seq). Each dataset included samples taken at discrete circadian time-points from wild-type mice and clock gene knockout mice (Suppl. Table S1). For each dataset, we used RAIN to detect genes rhythmic in at least one genotype, applying a less stringent cutoff (q_rhy_ ≤ 0.1) to have more genes for comparison. For rhythmic genes, we then used LimoRhyde and DODR to calculate q-values of differential rhythmicity (Fig. 2A). The median runtime of LimoRhyde and limma was 0.3 seconds per dataset, whereas the median runtime of DODR was 2 minutes per dataset.

**Figure 2.**
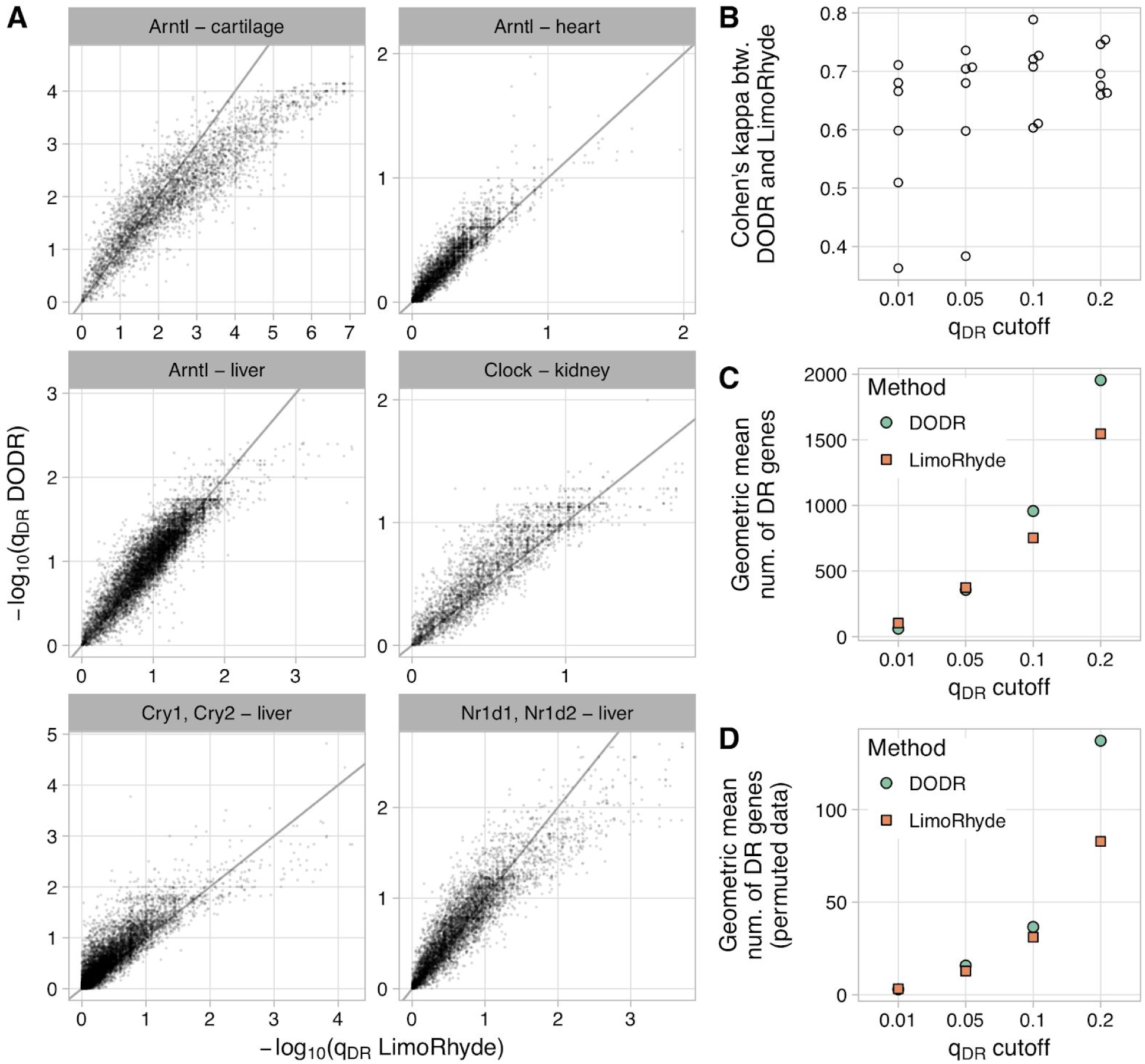
Comparing LimoRhyde (followed by limma) and DODR for detecting differential rhythmicity (DR) between wild-type and clock gene knockout mice. For details of datasets, see Suppl. Table S1. In each dataset, rhythmic genes were identified using RAIN (q_rhy_ ≤ 0.1). **(A)** Scatterplots of q-value of differentially rhythmicity as calculated by each method. The title of each plot indicates the knocked-out gene(s) and the tissue in which gene expression was measured. Each point represents a rhythmic gene. The line indicates y = x. For each dataset, up to 15 genes with extremely high −log_10_(q_DR_ LimoRhyde) are not shown. **(B)** Cohen’s kappa, a measure of inter-rater agreement, between DODR and LimoRhyde at various q-value cutoffs. Each point represents a dataset. **(C)** Geometric mean (across datasets) of the number of differentially rhythmic genes at various q-value cutoffs. **(D)** Geometric mean (across datasets) of the mean (across permutations) number of differentially rhythmic genes at various q_DR_ cutoffs, in data in which the sample labels (wild-type or knockout) were permuted. Labels were permuted after identifying rhythmic genes, and were only permuted within samples at the same time-point. Thus, DR genes identified in permuted data can be considered false positives for differential rhythmicity.

Overall, q-values from the two methods were highly correlated (median Pearson correlation 0.90). In addition, based on Cohen’s kappa, LimoRhyde and DODR showed moderate to strong agreement at various q-value cutoffs (Fig. 2B). Although the number of differentially rhythmic genes varied between datasets, LimoRhyde tended to select slightly more genes than DODR at a low q-value cutoff (q_DR_ ≤ 0.01) and somewhat fewer genes at higher q-value cutoffs (q_DR_ ≤ 0.1 or q_DR_ ≤ 0.2; Fig. 2C and Suppl. Fig. S2A). To evaluate the ability of the two methods to control false positives, we performed permutation testing on each dataset (see Materials and Methods). Both methods effectively controlled false positives, detecting many fewer differentially rhythmic genes on permuted data than on the unpermuted data, although again LimoRhyde tended to select fewer genes (i.e., was more conservative) than DODR at higher q-value cutoffs (Fig. 2D and Suppl. Fig. S2B). These results suggest that LimoRhyde (followed by limma) and DODR provide comparable detection of differential rhythmicity in circadian transcriptome data.

### Applying LimoRhyde to human transcriptome data from diverse experimental designs

To explore the flexibility of LimoRhyde, we used it to analyze two transcriptome datasets from humans, each of which has a different experimental design than the datasets from mice. The first dataset from humans was based on brain tissue from postmortem donors, with the zeitgeber time for each sample based on the respective donor’s geographic location, date, and time of death (Chen et al., 2016). The 146 donors ranged in age from 16 to 96 years old (50.7 ± 15.3, mean ± SD). Given how sleep-wake patterns change with age (Roenneberg et al., 2004; Yoon et al., 2003), this dataset presents an excellent opportunity to examine the interaction between aging and circadian rhythms in two regions of the human prefrontal cortex (Brodmann’s areas 11 and 47). The original analysis, however, was forced to discretize donors into younger and older, which discards information and sacrifices statistical power. LimoRhyde, on the other hand, can accommodate continuous variables such as age without discretizing them.

Because the time-points are based on times of death, they are approximately randomly spaced, precluding the use of RAIN or JTK_CYCLE. Therefore, to identify rhythmic genes, we used an additive model in LimoRhyde, including terms for age, zeitgeber time, and brain region (Fig. 3A; three example genes are shown in Fig. 3B-C). This additive model is equivalent to cosinor regression. To estimate each gene’s overall rhythm amplitude, we applied ZeitZeiger to the residuals of an additive model lacking terms for zeitgeber time (see Materials and Methods). Applying the criteria of q_rhy_ ≤ 0.1 and rhythm amplitude ≥ 0.1, we identified 891 genes as rhythmic (Fig. 3D and Suppl. Fig. S3A).

**Figure 3.**
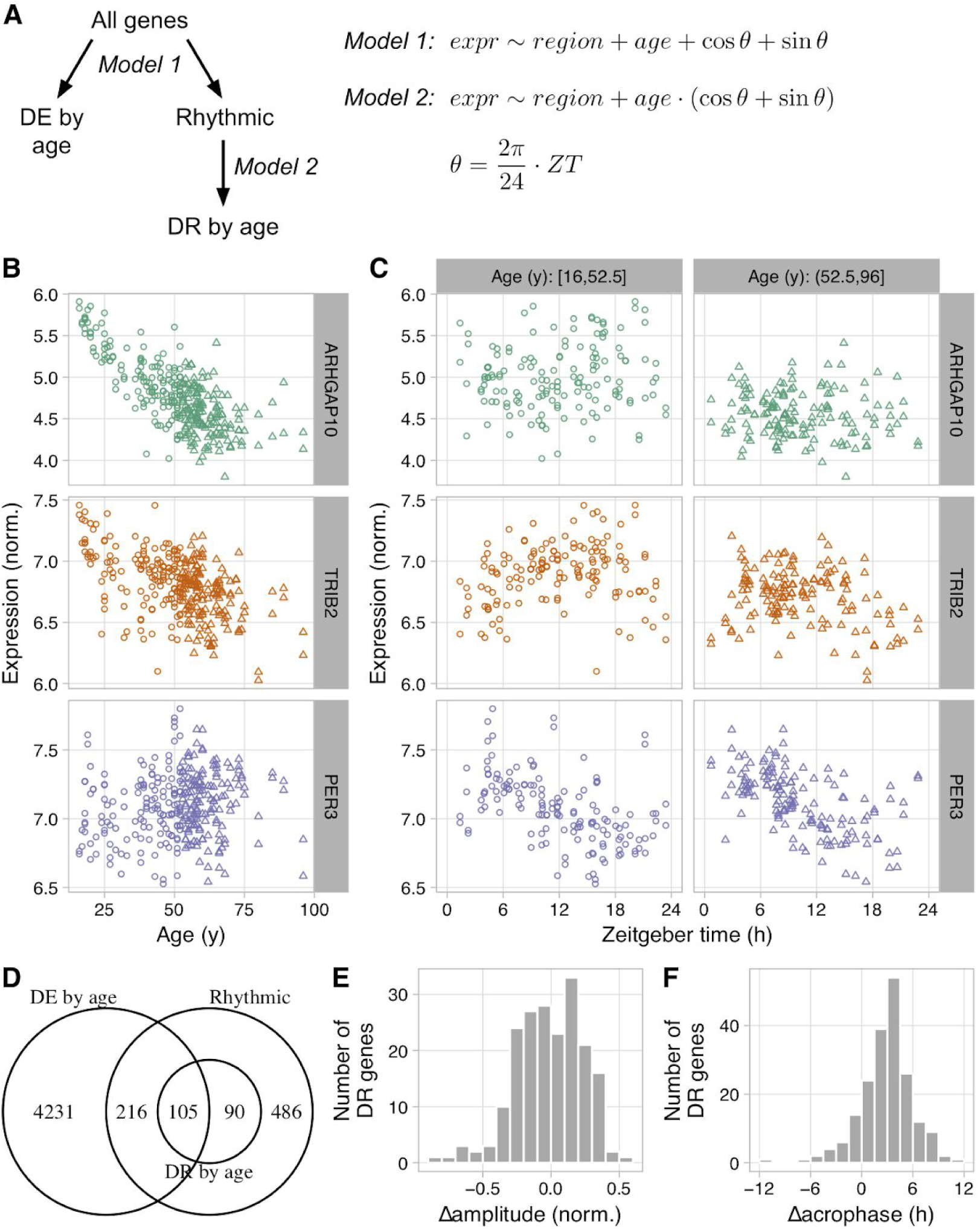
Using LimoRhyde to analyze transcriptome data based on postmortem samples from human brain (GSE71620). **(A)** Schematic of workflow and linear model formulae used to detect rhythmic, differentially rhythmic (DR), and differentially expressed (DE) genes. **(B)** and **(C)** Scatterplots for three example genes, showing log-normalized expression as a function of age and as a function of zeitgeber time of death within younger and older donors. Each point represents a sample. ARHGAP10 is classified as DE only, TRIB2 as DE and DR, and PER3 as rhythmic only. **(D)** Venn diagram of genes meeting criteria for rhythmicity (q_rhy_ ≤ 0.1 and rhythm amplitude ≥ 0.1), differential rhythmicity (q_DR_ ≤ 0.1), and differential expression (q_DE_ ≤ 0.01). **(E)** and **(F)** Histograms of Δamplitude and Δacrophase for differentially rhythmic genes between younger and older donors, calculated using ZeitZeiger. Positive Δamplitude indicates higher rhythm amplitude in older donors. Positive Δacrophase indicates a phase advance in older donors.

Using the original additive model, we then identified 4,551 genes whose expression increased or decreased with age, accounting for zeitgeber time (q_DE_ ≤ 0.01; Fig. 3D and Suppl. Fig. S3B). Genes whose expression decreased with age were strongly enriched for involvement in glutamate receptor signaling, synapse structure and activity, and mitochondria (Suppl. Table S4), which is consistent with previous findings (Lu et al., 2004).

To find genes whose rhythmic expression varied with age, we altered the linear model to include an interaction between age and zeitgeber time (Fig. 3A). Of the 891 genes that met our criteria for rhythmicity, 196 genes were differentially rhythmic (q_DR_ ≤ 0.1; Fig. 3D and Suppl. Fig. S3C). For each differentially rhythmic gene, we used ZeitZeiger to estimate rhythm amplitude and acrophase within the younger 50% and older 50% of donors. Changes in rhythm amplitude were centered near zero, with similar numbers of genes showing increased or decreased amplitude in older donors (Fig. 3E). Genes with decreased rhythm amplitude were enriched for involvement in leukocyte-mediated immunity and the adaptive immune response (Suppl. Table S5). Changes in acrophase were shifted from zero, corresponding to a mean advance of 3.1 h in older donors (circular mean; Fig. 3F).

The second dataset from humans was based on suction-blister epidermis samples acquired from 20 subjects at three time-points (9:30am, 2:30pm, and 7:30pm) (Spörl et al., 2012). The original analysis, which used limma but considered the time-points as categorical variables (a la ANOVA) and did not adjust for inter-subject variation, identified 294 genes whose expression varied with time of day (q ≤ 0.05).

To analyze the dataset using LimoRhyde, we constructed a linear model with terms for subject and time of day (Suppl. Fig. S4A). We then used limma to perform a moderated F-test on the two coefficients corresponding to time of day, which identified 1,436 genes with time-of-day-dependent expression (q ≤ 0.05; Suppl. Fig. S4B-C). Among the 15 top-ranked genes were eight core clock genes (NR1D1, PER3, CIART, NPAS2, PER1, ARNTL, NR1D1, and PER2, all with q ≤ 2·10^−8^).

Because this dataset has exactly three time-points, the LimoRhyde time decomposition and ANOVA are equivalent; they both correspond to two parameters in the linear model (the increased number of detected genes in our analysis is a result of adjusting for inter-subject variation). As the number of time-points increases, though, LimoRhyde will continue to favor genes whose expression varies sinusoidally over time, whereas ANOVA, which ignores the relationship between time-points, will not. Taken together, these examples demonstrate how LimoRhyde enables statistically rigorous analysis of circadian transcriptome data from diverse experimental designs.

## Discussion

Despite the increasingly complexity of experiments to interrogate rhythmic biological systems, methods to analyze the resulting genome-scale data have remained largely ad hoc. Here we described LimoRhyde, a unified approach to detect gene-wise differential rhythmicity and differential expression in circadian or otherwise rhythmic transcriptome data. LimoRhyde is inspired by cosinor regression and is applicable to data from any experimental design that can be described by a linear model. LimoRhyde thus functions as an adapter, making circadian transcriptome data amenable to analysis by the ever-improving and growing set of methods designed for differential analysis of microarray and RNA-seq data.

For detecting differential rhythmicity in the common two-condition scenario, our results suggest that LimoRhyde performs similarly to DODR. Although LimoRhyde (followed by limma) is considerably faster, the absolute difference in runtime is trivial compared to the amount of time required to perform the experiments. On some datasets, LimoRhyde followed by limma may be more conservative than DODR, implying that one could use a higher q-value cutoff to capture a similar number of differentially rhythmic genes at a similar false discovery rate.

LimoRhyde distinguishes itself by its versatility. First, LimoRhyde can be used to detect rhythmic or time-of-day-dependent gene expression in datasets in which time-points are either randomly spaced or do not cover the full circadian cycle, scenarios for which methods such as JTK_CYCLE and RAIN are ill-suited. In this application, LimoRhyde is conceptually equivalent to cosinor regression, with the advantage of using empirical Bayes procedures in methods such as limma to share information between genes. Second, LimoRhyde enables the detection of differential expression between conditions, accounting for possible rhythmicity. This could reveal expression changes in genes whose mRNAs are too stable to be rhythmic or differentially rhythmic (Lück et al., 2014). Our results suggest that a typical circadian experiment is well powered to detect even relatively small log fold-changes. Third, LimoRhyde can be applied to transcriptome data from complex experimental designs. Here we analyzed a dataset in which an experimental variable was continuous and a dataset in which multiple samples were collected from each participant.

While LimoRhyde provides rigorous p-values, other methods are useful for interpretation. For example, given a set of differentially rhythmic genes, methods such as ZeitZeiger can quantify the changes in rhythm amplitude and phase. Furthermore, gene set analysis methods such as CAMERA can identify biological processes that are enriched for changes in average expression level or in rhythm amplitude. An analogous method called Phase Set Enrichment Analysis could identify processes enriched for changes in phase (Zhang et al., 2016).

Regardless of the computational method, detection of differential rhythmicity and differential expression requires two assumptions. First, one must assume a value for the period of the rhythm. For typical experiments using entrained or free-running organisms, the assumed period should likely correspond to the period of the zeitgeber (T) or the free-running period of the organism (tau), respectively. Second, one must assume an alignment of the time-points between different conditions. For example, if samples are collected from organisms in different photoperiods, the results will depend on whether time 0 in each photoperiod is defined as the time of lights on or the time of lights off. Consequently, we advise caution when calculating and interpreting differential rhythmicity and differential expression in datasets based on free-running organisms for which tau varies considerably between conditions. The danger of this experimental design is that, if the time-points are not aligned properly, the results will be confounded by differences in the organisms’ intrinsic circadian phase (Hsu and Harmer, 2012). In addition to these assumptions, drawing a distinction between rhythmic, differentially rhythmic, and differentially expressed genes — although convenient — requires arbitrary cutoffs of q-value and/or rhythm amplitude. An alternative approach would be to test the coefficients for the main effect and the statistical interaction jointly, which would identify genes showing evidence for either differential rhythmicity or differential expression.

Multiple features of LimoRhyde remain to be explored. For example, although in this paper we used LimoRhyde in conjunction with limma, which is fast and can handle both microarray and RNA-seq data, LimoRhyde is compatible with multiple other methods for differential expression analysis. In addition, although here we decomposed time using sine and cosine curves (as in cosinor), it is also possible to apply a decomposition based on a periodic smoothing spline (as in ZeitZeiger). LimoRhyde could also be used to detect differences in higher-order harmonics of circadian gene expression (Hughes et al., 2009).

In conclusion, we have developed a general approach to analyze rhythmic transcriptome data in which there are multiple experimental variables. Here we concentrated on microarray and RNA-seq data, but given limma’s success on proteomics, DNA methylation, and ChIP-Seq data (Brusniak et al., 2008; Lun and Smyth, 2014; Maksimovic et al., 2012), we are optimistic that LimoRhyde could be applied to other types of genome-scale data as well. Altogether, LimoRhyde can help ensure that our ability to analyze rhythmic omics data continues to scale with our ability to acquire it.

## Acknowledgments

I thank Seth Rhoades for helpful comments on the manuscript. This work was supported by NIH R35 GM124685 to J.J.H.

## Conflict of Interest Statement

The author declares that there is no conflict of interest.

