## Supplementary Materials for "LimoRhyde: a flexible approach for differential analysis of rhythmic transcriptome data"

Jacob J. Hughey[1,2,*]

[1]Department of Biomedical Informatics, Vanderbilt University School of Medicine, Nashville, Tennessee; [2]Department of Biological Sciences, Vanderbilt University, Nashville, Tennessee

Figure S1

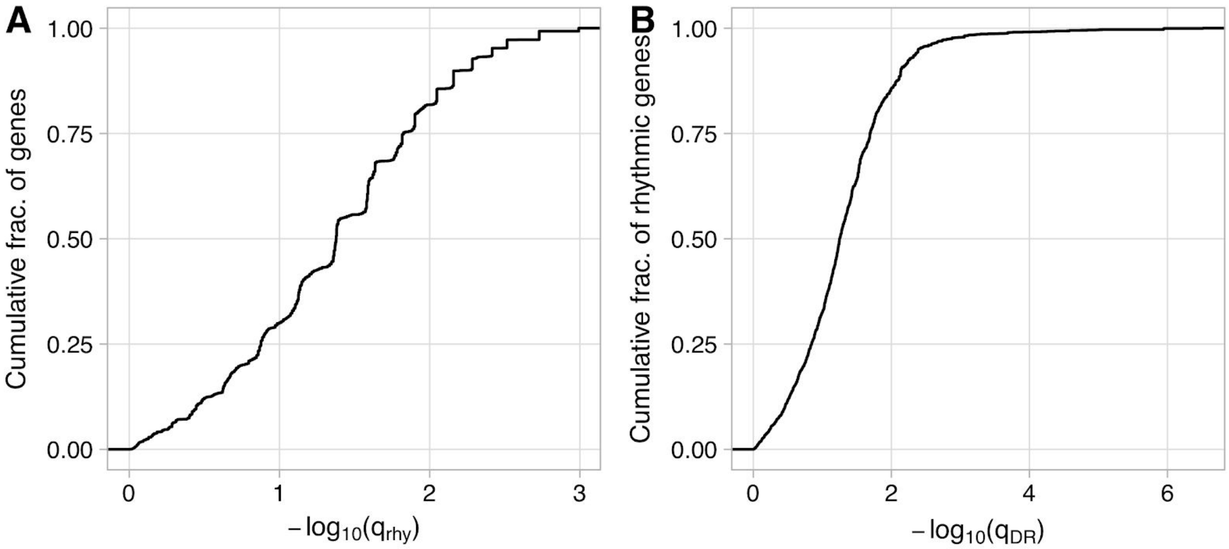


Analyzing circadian transcriptome data from liver of wild-type and Arntl-/- mice under night-restricted feeding (GSE73552). **(A)** Empirical cumulative distribution function for $-\log_{10}(q_{rhy})$, where $q_{rhy}$ corresponds to a gene's q-value of being rhythmic in at least one genotype, calculated using RAIN. **(B)** Empirical cumulative distribution function for $-\log_{10}(q_{DR})$, where $q_{DR}$ corresponds to a rhythmic gene's q-value of differential rhythmicity, calculated using LimoRhyde and limma and based only on genes for which $q_{rhy} \leq 0.01$.

Figure S2

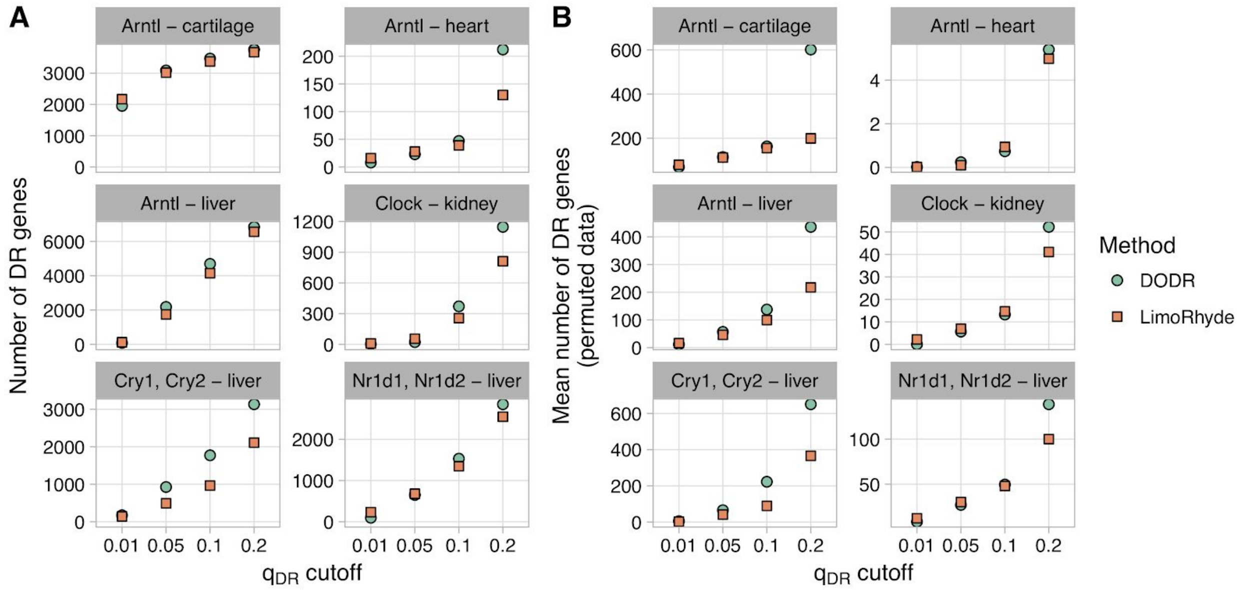


Comparing LimoRhyde (followed by limma) and DODR for detecting differential rhythmicity between wild-type and clock gene knockout mice. The title of each plot indicates the knocked-out gene(s) and the tissue in which gene expression was measured. For details of datasets, see Suppl. Table S1. In each dataset, rhythmic genes were identified using RAIN ($q_{rhy} \leq 0.1$). **(A)** Number of differentially rhythmic (DR) genes at various q-value cutoffs. **(B)** Number of differentially rhythmic genes at various q-value cutoffs, in data in which the sample labels (wild-type or knockout) were permuted. Labels were permuted after identifying rhythmic genes, and were only permuted within samples at the same time-point. Thus, DR genes identified in permuted data can be considered false positives for differential rhythmicity.

Figure S3

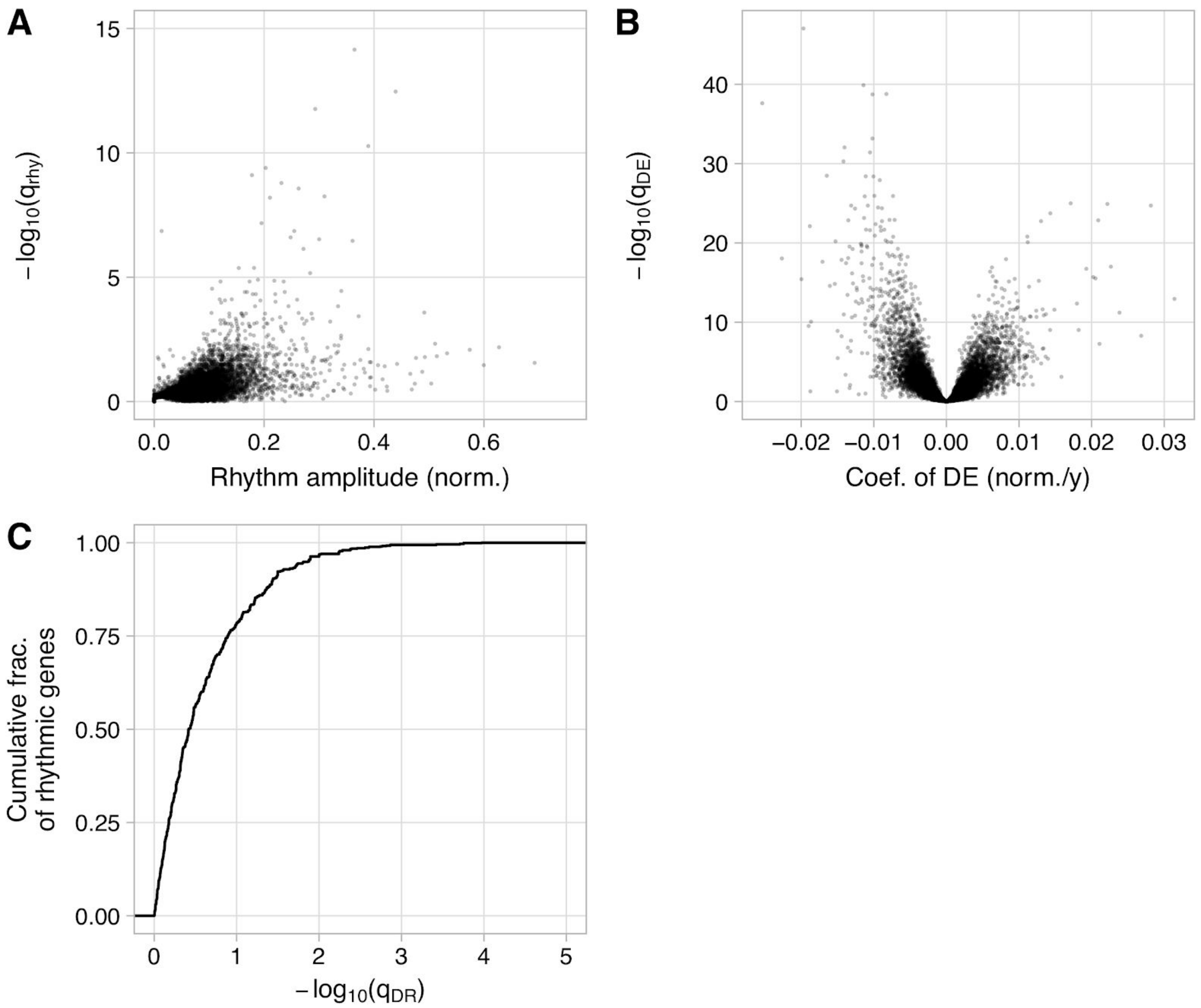


Analyzing circadian transcriptome data based on postmortem samples from human brain (GSE71620). **(A)** Scatterplot of -$\log_{10}(q_{rhy})$ vs. rhythm amplitude, where $q_{rhy}$ is the q-value of rhythmicity. Each point represents a gene. $q_{rhy}$ was calculated using LimoRhyde and limma with an additive model including terms for zeitgeber time, age, and brain region. Rhythm amplitude was calculated using ZeitZeiger and the residuals of an additive model including terms for age and brain region. Rhythm amplitude is in log-normalized units of expression. **(B)** Scatterplot of -$\log_{10}(q_{DE})$ vs. the coefficient of differential expression, where $q_{DE}$ is the q-value of differential expression. Each point represents a gene. Because age is continuous, the coefficient does not correspond to a log fold-change. **(C)** Empirical cumulative distribution function of -$\log_{10}(q_{DR})$, where $q_{DR}$ is the q-value of differential rhythmicity, based on a linear model with an interaction age and zeitgeber time, considering only genes having $q_{rhy} \leq 0.1$ and rhythm amplitude $\geq 0.1$.

Figure S4

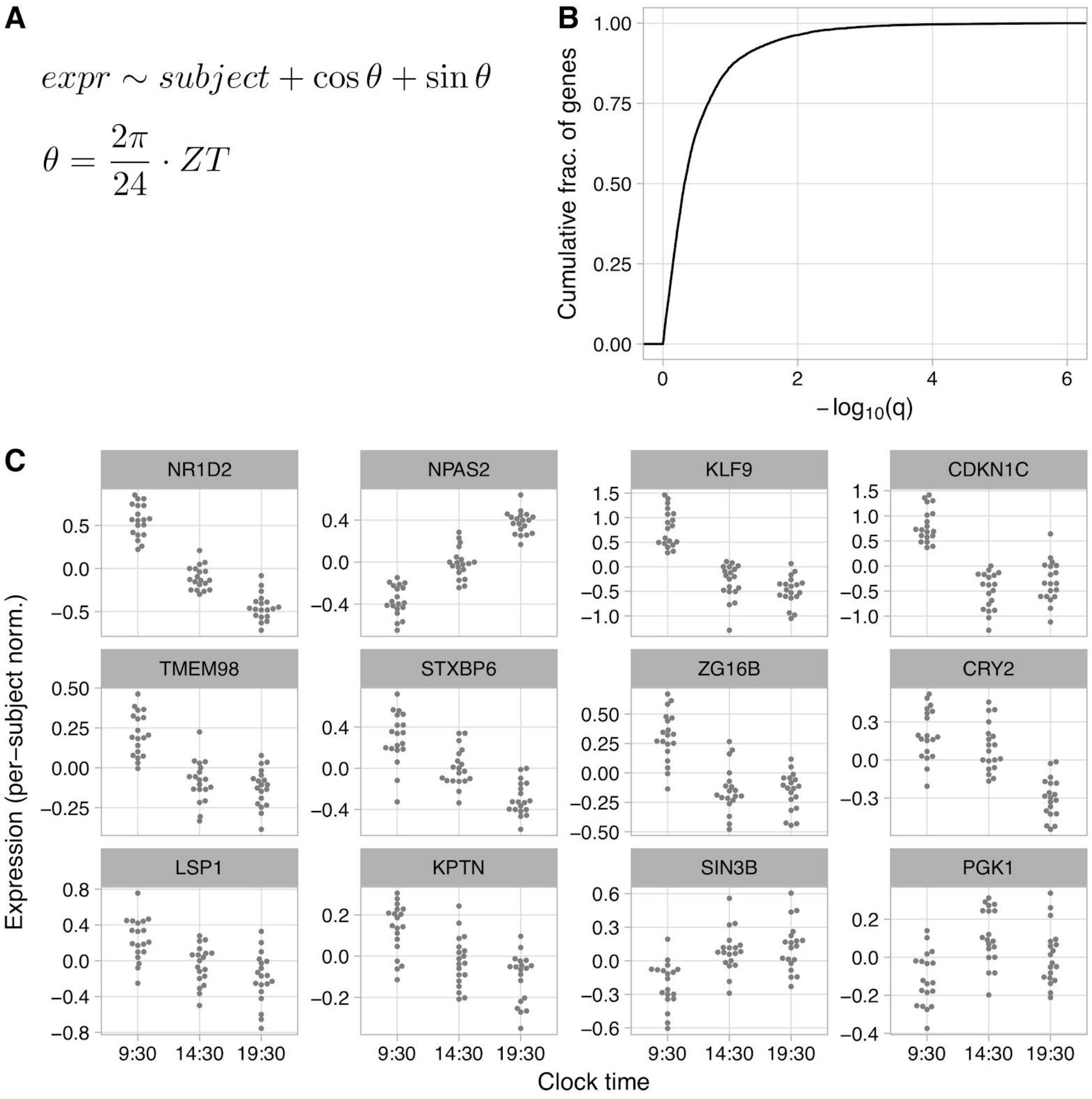


Using LimoRhyde identify genes whose expression varies with time of day in human epidermis (GSE35635). **(A)** Linear model formula passed to limma. **(B)** Empirical cumulative distribution function of q-value of time-of-day-dependent expression. Twenty-seven genes with $q \leq 10^{-6}$ are not shown. **(C)** Per-subject normalized expression for 12 example genes. Each point represents a sample. Expression values correspond to the residuals of a limma fit based only on subject. Genes in the top row have $q \leq 10^{-8}$, genes in the middle row have $10^{-6} < q \leq 10^{-4}$, and genes in the bottom row have $10^{-3} < q \leq 0.1$.
